# Conservation of conformational dynamics across prokaryotic actins

**DOI:** 10.1101/260208

**Authors:** Natalie Ng, Handuo Shi, Alexandre Colavin, Kerwyn Casey Huang

## Abstract

The actin family of cytoskeletal proteins is essential to the physiology of virtually all archaea, bacteria, and eukaryotes. While X-ray crystallography and electron microscopy have revealed structural homologies among actin-family proteins, these techniques cannot probe molecular-scale conformational dynamics. Here, we use all-atom molecular dynamic simulations to reveal conserved dynamical behaviors in four prokaryotic actin homologs: MreB, FtsA, ParM, and crenactin. We demonstrate that the majority of the conformational dynamics of prokaryotic actins can be explained by treating the four subdomains as rigid bodies. MreB, ParM, and FtsA monomers exhibited nucleotide-dependent dihedral and opening angles, while crenactin monomer dynamics were nucleotide-independent. We further determine that the opening angle of ParM is sensitive to a specific interaction between subdomains. Steered molecular dynamics simulations of MreB, FtsA, and crenactin dimers revealed that changes in subunit dihedral angle lead to intersubunit bending or twist, suggesting a conserved mechanism for regulating filament structure. Taken together, our results provide molecular-scale insights into the nucleotide and polymerization dependencies of the structure of prokaryotic actins, suggesting mechanisms for how these structural features are linked to their diverse functions.

**Significance Statement:** Simulations are a critical tool for uncovering the molecular mechanisms underlying biological form and function. Here, we use molecular-dynamics simulations to identify common and specific dynamical behaviors in four prokaryotic homologs of actin, a cytoskeletal protein that plays important roles in cellular structure and division in eukaryotes. Dihedral angles and opening angles in monomers of bacterial MreB, FtsA, and ParM were all sensitive to whether the subunit was bound to ATP or ADP, unlike in the archaeal homolog crenactin. In simulations of MreB, FtsA, and crenactin dimers, changes in subunit dihedral angle led to bending or twisting in filaments of these proteins, suggesting a mechanism for regulating the properties of large filaments. Taken together, our simulations set the stage for understanding and exploiting structure-function relationships of bacterial cytoskeletons.

## Introduction

The eukaryotic cytoskeleton, which is critical for many cellular functions such as cargo transport and morphogenesis, is comprised of three major elements: actin, tubulin, and intermediate filaments. These proteins bind nucleotides and form highly dynamic polymers [1]. Each of these proteins has numerous homologs across the bacterial and archaeal kingdoms that dictate cell shape and various intracellular behaviors [1, 2]. However, relatively little is known about the structural dynamics of these prokaryotic homologs and whether dynamical behaviors are conserved.

Among bacterial cytoskeletal proteins, actin homologs are the most structurally and functionally diverse class identified thus far. Although sequence homology to eukaryotic actin is generally low, prokaryotic actins have been identified via X-ray crystallography based on their structural homology to eukaryotic actin [3–6], which has a U-shaped four-domain substructure, with two beta domains and a nucleotide binding pocket between two alpha domains [7]. Among the actin homologs, one of the best studied is MreB, which forms filaments that coordinate cell-wall synthesis in many rodshaped bacteria and is essential for maintaining cell shape in these species [8, 9]. FtsA is an actin homolog with a unique structural domain swap that is essential for anchoring the key cell-division protein and tubulin homolog FtsZ to the membrane [5, 10]. The actin homolog ParM forms filaments that move R1 plasmids to opposite ends of rodshaped bacteria prior to cytokinesis [11]. Crenactin forms part of the archaeal cytoskeleton [12]; its biological function is currently unknown, but is hypothesized to be involved in DNA segregation and/or cell-shape control [12]. Given the common structural features of prokaryotic actins, it is unknown how they exert such a wide variety of functions. Features such as the domain swap in FtsA suggest that some proteins may have the capacity for unique intramonomeric conformational changes [13]. Another possibility is that functional differences emerge at the filament level: a wide variety of double-protofilament bacterial-actin filament structures have been observed [14, 15]. The extent to which lessons about structure-function relationships are general across the diverse actin family can be informed by understanding commonalities and distinctions in their structural dynamics.

While X-ray crystallography and cryo-electron microscopy (cryo-EM) have proven critical for elucidating the structures of monomers and filaments of prokaryotic actins, understanding the mechanisms by which these proteins exert their functions, particularly their mechanical roles, requires integration with other experimental and computational techniques. Microscopy has revealed that most actin homologs can form long filaments within cells [3, 4, 16–19]. *In vitro*, ParM filaments exhibit dynamic instability [20], and all actin homologs except FtsA have been observed to undergo nucleotide hydrolysis [12, 21, 22]. However, these experimental techniques lack the spatial and temporal resolution necessary to understand how these filament properties are linked to changes in structure.

Various mechanistic models of cytoskeletal function have focused on nucleotide hydrolysis as a key determinant of filament mechanics [23–25]. Understanding how nucleotide hydrolysis and polymerization affect structural transitions in prokaryotic actins requires a method that can interrogate molecular behaviors with atomic resolution. All-atom molecular dynamics (MD) simulations have been successfully employed to probe the effects of perturbations on prokaryotic and eukaryotic cytoskeletal proteins. MD simulations of eukaryotic actin monomers have uncovered nucleotide-dependent changes in the structure of the nucleotide-binding pocket [26], and simulations of actin filaments showed nucleotide-dependent changes to filament bending [27]. MD simulations predicted that GTP hydrolysis of the tubulin homolog FtsZ can result in substantial polymer bending [28], which was subsequently verified through X-ray crystallography [29]. MD simulations of MreB and FtsA filaments also revealed intra- and inter-subunit changes with important implications for their respective cellular functions [13, 17]. In sum, structural changes to cytoskeletal filaments are generally observable within the time frame accessible to MD simulations, potentiating a systematic survey of general and specific connections among bound nucleotide, polymerization, and subunit conformations across the prokaryotic actin family.

Here, we used MD simulations to probe the conformational dynamics of monomers and filaments of MreB, FtsA, ParM, and crenactin (Fig. 1). We found that these proteins exhibit a wide range of intrasubunit motions that are generally well described by the centers-of-mass of their four subdomains, and hence the majority of monomer dynamics can be explained by changes in opening and dihedral angles formed by the subdomain centers. Our results predict that some proteins exhibit strong dependence on the bound nucleotide, while others are unaffected by hydrolysis. In ParM, opening is inhibited by interactions between two subdomains. As with MreB, changes in the dihedral angle of FtsA and crenactin subunits generally impact the bending or twisting of polymers. This work provides insight into how molecular-scale perturbations of these proteins contribute to their diverse roles in cell-shape regulation and intracellular organization across bacteria and archaea.

**Figure 1:**
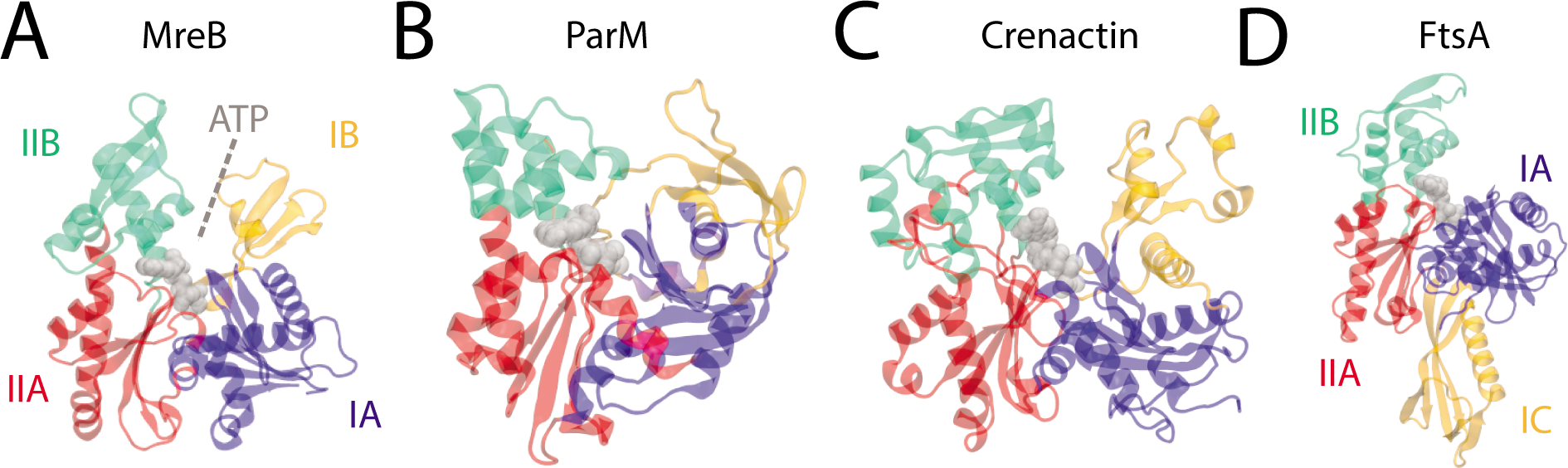
Structures of prokaryotic actin homologs. A-C) The crystal structures of (A) MreB (PDB ID: 1JCG), (B) ParM (PDB ID: 1MWM), and (C) crenactin (PDB ID: 4CJ7) display a characteristic U-shaped actin-like fold described by four subdomains surrounding an enclosed ATP-binding pocket (gray). D) The crystal structure of FtsA (PDB ID: 4A2B) shows a domain swap of IB to IC.

## Results

### The nucleotide-dependent conformational dynamics of MreB are well represented by the centers of four subdomains

In a previous study, we performed all-atom MD simulations on unconstrained MreB monomers using CHARMM27 force fields and found that ATP-bound monomers had larger opening and dihedral angle than ADP-bound monomers [13]. For our study of prokaryotic actins, we first sought to interrogate the robustness of these findings with respect to the force field used and the dimensional reduction to the centers-of-mass of subdomains IA, IB, IIA, and IIB of actin family members.

While simulations using different force fields mostly preserve large-scale motions, distinct behaviors emerge at finer levels of detail [30]. Thus, we performed allatom MD simulations on *Thermatoga maritima* MreB (PDB ID: 1JCG) [4] using CHARMM36 force fields [31]. As done previously for actin [32] and MreB [28], we quantified conformational changes by calculating two opening angles and a dihedral angle from the center of mass of each of the four subdomains (Methods). While the opening angle was 5-10° smaller with CHARMM36 than with CHARMM27 [13] (Fig. 2A,B, S1A), in both sets of simulations subdomains IB and IIB of ATP-bound monomers rapidly hinged apart to form stable, open conformations. Additionally, using CHARMM36, the opening angle equilibrated at smaller angles for ADP- than ATP- bound MreB (Fig. S1B), as expected from our previous study [13]. ATP-bound MreB monomers also adopted a larger dihedral angle than ADP-bound monomers using CHARMM36, similar to CHARMM27 (Fig. 2C,D, Fig. S1C). Thus, despite small differences, a similar nucleotide dependence in the conformation of MreB monomers was observed using both CHARMM27 and CHARMM36 force fields, supporting our use of CHARMM36 going forward.

**Figure 2:**
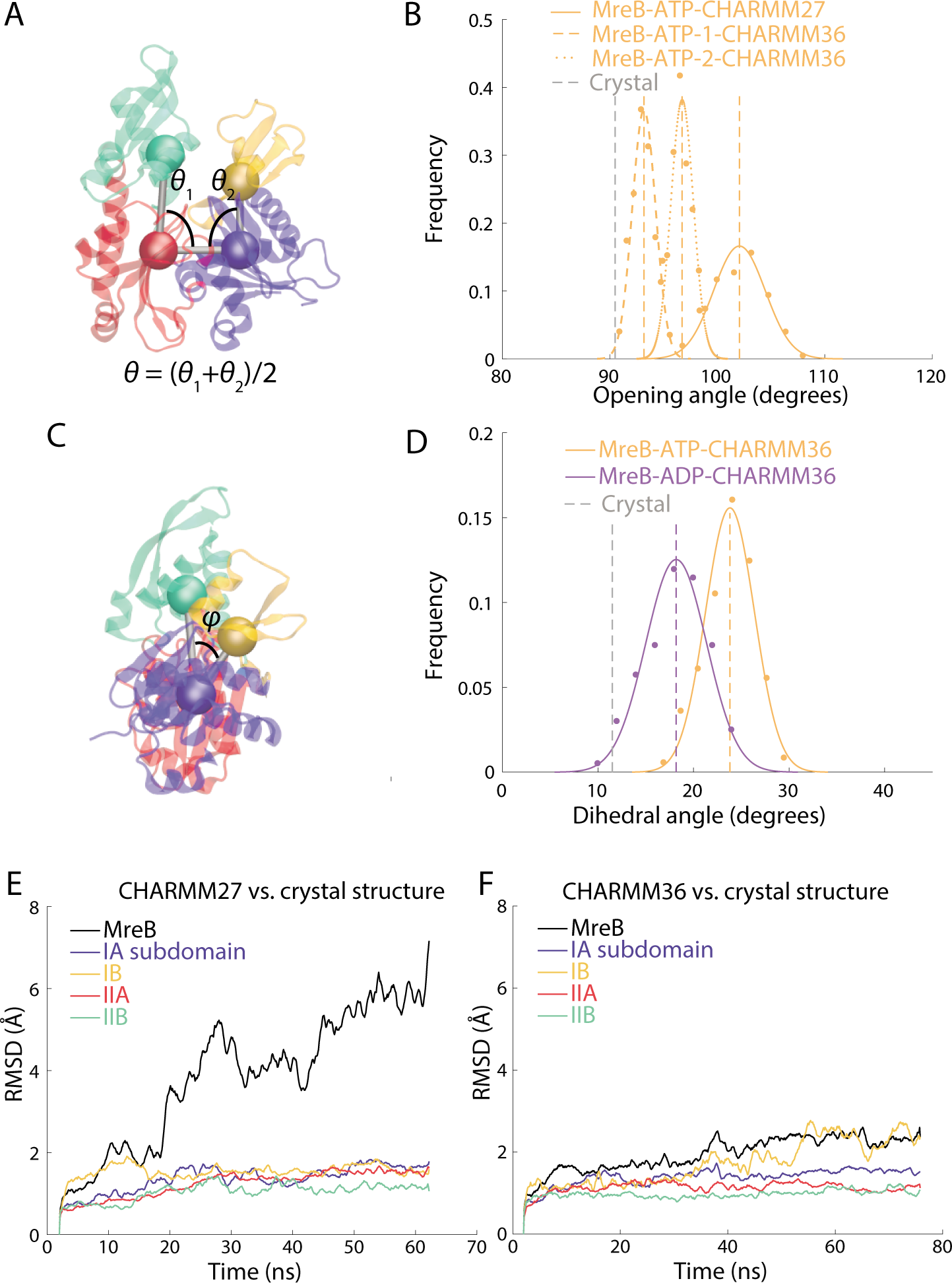
MreB adopts multiple conformations with nucleotide-dependent opening and dihedral angles. A) The opening angle of an MreB monomer is defined as the average of the internal opening angles. B) The opening angle distribution in the last 30 ns of simulation is larger for ATP- bound than ATP-bound MreB monomers. The opening angle of an ATP-bound MreB monomer equilibrated at an even larger value in a CHARMM27 simulation. The rest of the simulations in this manuscript use CHARMM36 force fields, unless otherwise noted. Dashed lines are mean values. Gray dashed line is value in crystal structure. C) Schematic illustrating calculation of the dihedral angle. D) Histograms of the dihedral angle during the last 30 ns of the simulations show that an ATP-bound MreB monomer adopts a larger dihedral angle than an ADP- bound MreB monomer. Dashed lines are mean values. Gray dashed line is value in crystal structure. E) The trajectory of the RMSD values of the MreB-ATP monomer in a CHARMM27 simulation relative to the initial equilibrated structure exhibited large changes as the protein adopted an open conformation (black line). Nonetheless, the RMSDs of the four subdomains remained ~2 Å, indicating that conformational dynamics were small within each subdomain. F) The RMSD of the entire protein computed from the trajectory of for the CHARMM36 MreB-ATP-1 simulation relative to the initial equilibrated structure remained relatively low compared with (E). The RMSDs of the four subdomains remained ~2 Å.

While previous studies used the centers-of-mass of the four subdomains of actin-family proteins to dramatically reduce the dimensionality of the protein structure [4, 13, 32], it is also possible for conformational changes to arise within subdomains in addition to the hinges between them. To distinguish between these scenarios, we calculated the root mean square deviation (RMSD) of the *C*_*α*_ atoms from the energetically minimized structure for each subdomain separately, and also for the entire protein, at each time point of our simulations.

In the CHARMM27 ATP-bound simulation, the RMSD of the entire protein increased past 5 Å as the opening angle increased. However, the RMSD of each subdomain remained at ~2 Å (Fig. 2E), suggesting that most conformational changes were inter-subdomain. Unsurprisingly, since the CHARMM36 simulation adopted a smaller opening angle than the CHARMM27 simulation, the RMSD of the protein was smaller as well (Fig. 2F). Nonetheless, consistent with the CHARMM27 simulation, the RMSD of each subdomain was smaller than the RMSD of the whole protein (Fig. 2F). To determine whether subdomain structure was consistent between distinct MreB monomer conformations, we computed RMSDs between the CHARMM36 equilibrium structure and the CHARMM27 simulation at each time point. Since the CHARMM27 simulation adopted a larger opening angle than the CHARMM36 simulation, the RMSD of the whole protein increased relative to the CHARMM36 equilibrium structure. Still, the subdomain RMSDs remained at ~2 Å (Fig. S1D). Thus, the structure of each subdomain is largely maintained as the whole protein undergoes large conformational changes.

### FtsA monomers exhibit nucleotide-dependent conformational changes

We next investigated FtsA (PDB ID: 4A2B), an essential protein involved in tethering the key division protein FtsZ to the membrane [5, 10]. FtsA has a four-subdomain architecture similar to those of actin and MreB, but subdomain IB is replaced by a new subdomain (IC) located on the opposite side of subdomain IA (Fig. 1), that has no structural similarity to the actin subdomains [5]. To determine whether this domain swap impacts the conformational dynamics around the nucleotide-binding pocket and alters the coupling of dihedral/opening angles to nucleotide hydrolysis, we first carried out all-atom unconstrained MD simulations on ATP- and ADP-bound FtsA monomers.

While FtsA monomers showed little conformational flexibility, they still exhibited distinct ATP- and ADP-bound states with respect to opening and dihedral angles (Figure 3A,B, Methods). In all simulations, the RMSD of each subdomain as well as the entire protein remained <2 Å (Fig. S2), and the opening angle exhibited very little variation. Compared to an ATP-bound MreB monomer, whose opening angle reached a different equilibrium (102.1±2.4° and 93.2±1.0°, mean ± standard deviation (s.d.) measured over the final 30 ns of simulation) in replicate simulations, the opening angle of an ATP-bound FtsA monomer was much more constrained (109.7±0.8° and 109.6±0.8° in two replicates) and was highly reproducible (Fig. 3A,C,D). The FtsA equilibrium opening angle exhibited slight, but highly reproducible, nucleotide dependence: the opening angle for ADP-bound FtsA equilibrated at 111.8±0.7° and 111.5±0.7°. In ATP- and ADP-bound FtsA, the dihedral angle equilibrated at 20.6±1.9° for ATP and 20.3±2.5° for ADP, respectively (Fig. 3E,F), with a highly reproducible mean value across simulations (Fig. 3F). Thus, as with MreB and actin, FtsA likely has two distinct states dependent on the bound nucleotide.

**Figure 3:**
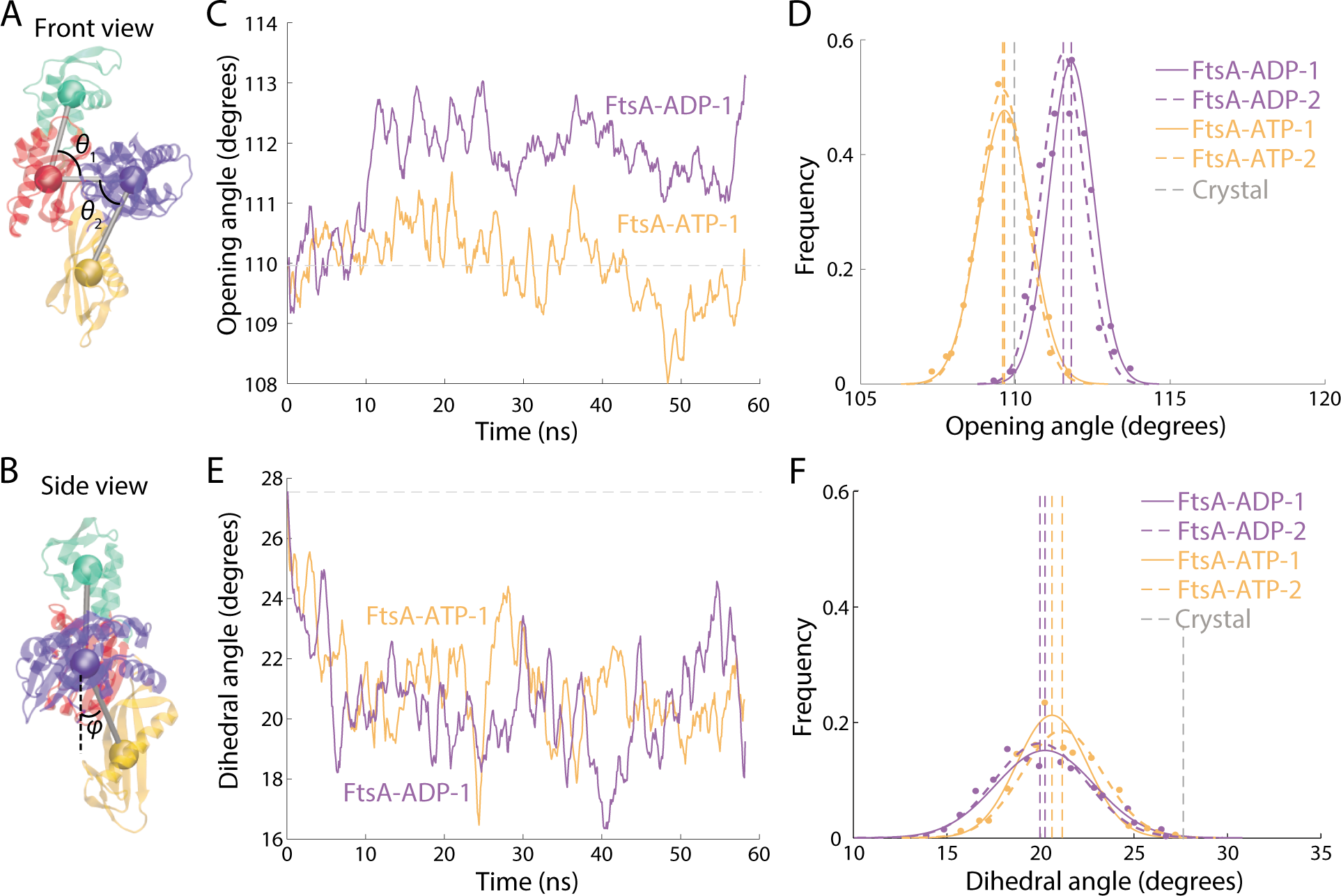
FtsA monomers undergo small but reproducible changes in opening angle upon nucleotide hydrolysis. A) The domain swap of IB to IC in FtsA necessitated a change in the calculation methodology for opening angle (Methods). B) Schematic of calculation methodology for FtsA dihedral angle. C) The opening angle of an ATP-bound FtsA monomer remained centered on the value in the crystal structure (gray dashed line), while an ADP-bound FtsA monomer equilibrated at a slightly larger opening angle. D) The distributions of opening angles over the last 30 ns of simulation were highly reproducible across the two replicate simulations for ATP- and ADP-bound FtsA monomers. Dashed lines are mean values. Gray dashed line is the value in the crystal structure. E,F) The trajectories (E) and distributions (F) of dihedral angles of ATP- and ADP- bound FtsA monomers were similar. Dashed lines are mean values. Gray dashed line is the value in the crystal structure.

### ParM exhibits high conformational variability with nucleotide-dependent states

We next used all-atom MD simulations to investigate ParM, which forms filaments that push apart plasmids to segregate them into daughter cells [6, 18]. Like MreB, ParM monomers exhibited large, nucleotide-dependent conformational changes, with substantial variability across replicate simulations. In all simulations of ATP-bound ParM, the opening angle rapidly increased from 97° in the crystal structure to over 100° (Fig. 4A). In one simulation, subdomains IB and IIB continued to hinge apart to 109.0±2.0° after 100 ns. In the other two simulations, the opening angle equilibrated at 102.2±1.4° and 102.2±1.7° ADP-bound monomers were less open, equilibrating between 97° and 99° (Fig. 4A). Unlike MreB, we did not observe consistent nucleotide dependencies on the dihedral angle of ParM monomers (Fig. S3).

**Figure 4:**
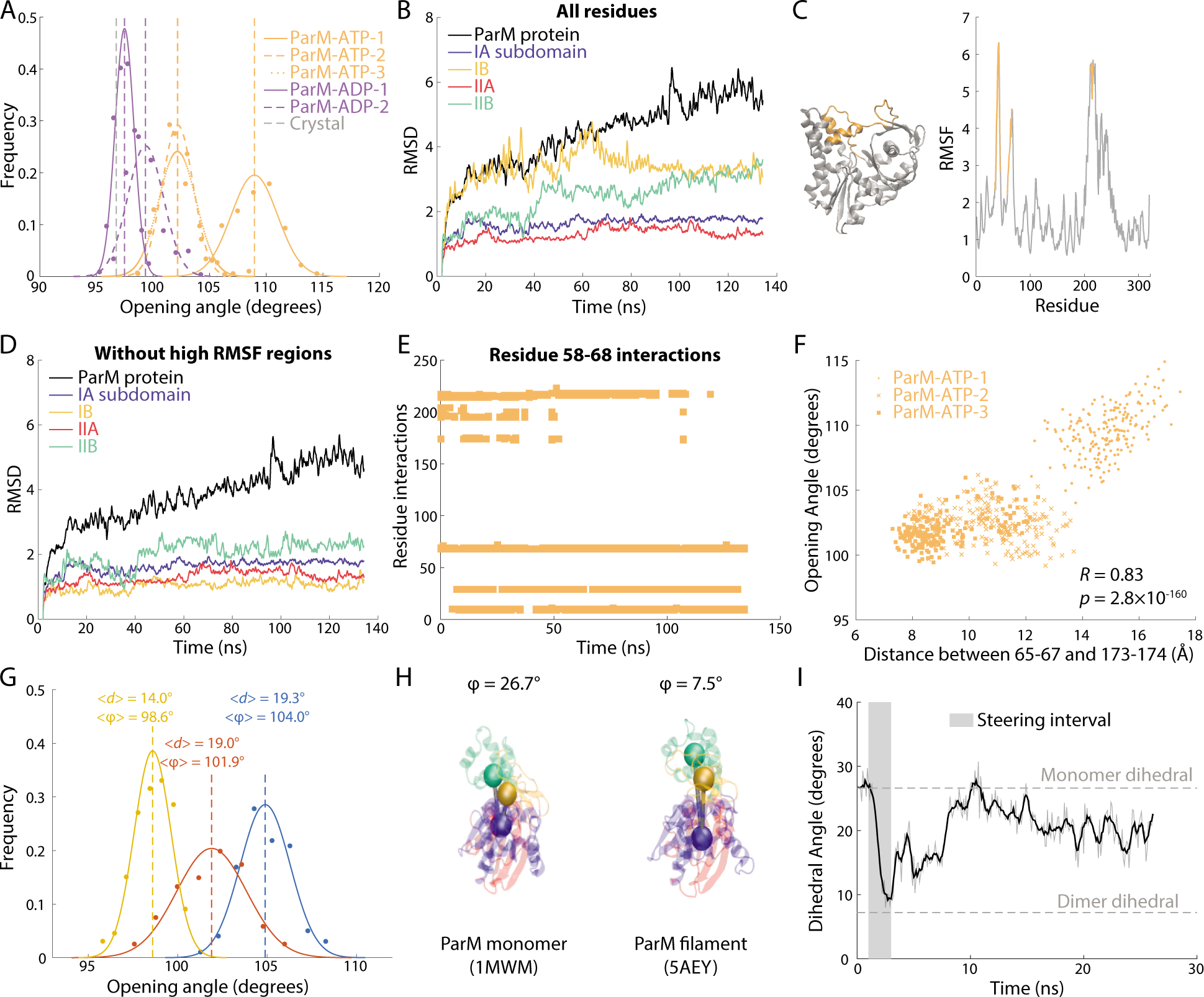
Loop in the IB domain drives ParM monomer opening. A) The opening angle of ADP-bound ParM monomers remained near the crystal structure value (gray dashed line), while that of ATP-bound ParM monomers consistently increased, with two simulations equilibrating between 100° and 104° and another opening beyond 105°. Dashed lines are mean values. Gray dashed line is the value in the crystal structure. B) For the ATP-bound ParM simulation in which the opening angle increased beyond 105° (ParM-ATP-1), there were large increases in RMSD across the entire protein and in subdomains IIA and IIB. C) RMSF analysis of single residue fluctuations during the simulation in (B) revealed two regions (residues 58-67 and 173-174, gold) with high RMSF values that were spatially proximal on the crystal structure. D) For the simulation in (B), when ignoring residues 58-67 and 173-174, the RMSDs of all four subdomains dropped to ~2 Å. Thus, these regions were responsible for the conformational variability in (B). E) Interactions of the loop formed by residues 58-67 with residues 173-174 and 200202 (blue boxes) disappeared early in simulation ParM-ATP-1. Interactions were defined as a minimum distance between residues of <5 Å. F) The distance *(d)* between residues 58-67 and 173-174 was highly correlated with the opening angle (j) across all simulations of ParM-ATP monomers. G) Steering of the distance between residues 58-67 and 173-174 tuned the opening angle in a distance-dependent manner. Dashed lines are mean values. H) The dihedral angle of a ParM monomer crystal structure (PDB ID: 1MWM) was much higher than that of each subunit in a ParM filament crystal structure (PDB ID: 5AEY). I) When the dihedral angle of a ParM-ATP monomer was steered to 7.5° (gray box) and then released, the angle re-equilibrated at a value similar to unconstrained simulations (Fig. 4A), indicating that ParM flattens upon polymerization.

In order to identify whether certain parts of ParM contributed to an opening angle of >105° in one of the ATP-bound simulations, we calculated the RMSD of each subdomain and the whole protein relative to the minimized structure in that simulation. Subdomains IB and IIB exhibited large conformational variability, similar to the protein as a whole (Fig. 4B). We identified residues 35-45 and residues 58-67 on subdomain IB and residues 211-216 on subdomain IIB as having the greatest root mean square fluctuation (RMSF) (Fig. 4C), a measure of the positional variability of specific residues. The subdomain RMSDs calculated after removing these high-RMSF residues decreased to <2 Å, suggesting a stable core within each subdomain of ParM (Fig. 4D). We re-measured opening and dihedral angles excluding these high-RMSF residues, and found that while the initial values changed, the same nucleotide dependencies relating to dihedral and opening angle were observed (Fig. S4).

The high degree of variability in opening angle across replicate simulations suggested the opportunity to identify the structural elements that underlie ParM opening. In the crystal structure, the high RMSF loop of residues 58-67 interact strongly (defined as a Ca-Ca distance <5 Å) with residues 173-174, which lie near the ATP binding pocket, as well as with residues 200-202 (Fig. 4E). In the ParM-ATP simulation with the largest opening angle, these interactions were largely abolished within 40 ns (Fig. 4E). By contrast, in the other two ParM-ATP simulations with smaller opening angles, the interaction between residues 58-67 and 173-174 persisted throughout the simulation (Fig. S5A,B). In one of these simulations, the interaction between residues 58-67 and 173-174 was initially disrupted but quickly recovered (Fig. S5A), consistent with the smaller increase in opening angle in this simulation. Across these three simulations, the opening angle was highly correlated with the distance between the center of mass of residues 65-67 and the center of mass of 173-174 (Fig. 4F).

To determine whether disrupting the interaction between residues 173-174 and 58-67 would cause ParM to open, we steered the center-of-mass distance between residues 173-174 and 65-67 from the crystal structure value of 9.3 Å to various larger values. In a steered simulation in which we steered the distance between residues 173-174 and 65-67 to 19.3±1.0 Å, the opening angle increased to 104.0±1.4° (Fig. 4G), suggesting that breaking this interaction directly changes the ParM protein conformation. Steering the distance between residues 173-174 and 65-67 (Fig. 4G) to 19.1±0.8 Å and 14.0±1.1 Å resulted in opening angles of 101.9±2.0° and 98.6±1.0°, respectively, indicating that the distance between residues 173-174 and 65-67 tunes the opening angle of ParM monomers.

The dihedral angles of ParM in a monomer crystal structure [6] and in a cryo-EM filament structure [18] were 26.7° and 7.54°, respectively (Fig. 4H), suggesting that polymerization impacts ParM conformations. ParM forms left-handed double-helical filaments that make MD simulations infeasible due to the large number of subunits required to mimic a biologically relevant system. To overcome this challenge and to glean information about whether ParM filaments flatten upon polymerization, we steered the dihedral angle of an ATP-bound ParM monomer to 7° to match that of the cryo-EM filament structure. Upon release, the monomer rapidly unflattened to 20° (Fig. 4I), suggesting that ParM monomers, like MreB [13], flatten upon polymerization. Thus, ParM likely has some similar conformational properties as MreB, even though the interactions between the flexible regions of subdomains IB and IIB unique to ParM provide tunability to its opening angle.

### The dihedral angle of prokaryotic actins is coupled to filament bending and twisting

For MreB, we previously discovered that the dihedral angle of the bottom subunit in a dimer simulation was directly coupled to dimer bending [13]. In particular, the intersubunit bending of ATP-bound MreB was correlated to the dihedral angle throughout each simulation, and steering the dihedral angle to a flatter conformation reduced the bending of a dimer structure [13]. We confirmed these findings for the CHARMM36 force field by steering the dihedral angle of the bottom subunit of an

MreB-ATP dimer to 23.1°, 28.3°, and 33.0°, and observed the expected inverse relationship between dihedral angle and filament bending (Fig. S6). Given the similarities between the dynamics of MreB and other bacterial actin homologs at the monomeric level, we hypothesized that other actin-like filaments may also exhibit intersubunit behaviors coupled to intrasubunit changes.

We performed MD simulations of dimers of FtsA (PDB ID: 4A2B) and *Pyrobaculum calidifontis* crenactin (PDB ID: 4CJ7); crenactin is an archaeal actin homolog for which our MD simulations of ATP- and ADP-bound monomers exhibited similar conformations (Fig. S7A,B). Dimer structures were initialized from repeated subunits of the appropriate crystal structure. Due to ParM’s complicated filament structure, which requires four points of contact per monomer, we were unable to construct biologically relevant ParM dimers with a stable interface *in silico* [33]. For each time step of dimer simulations, we measured two bending angles and one twisting angle between the subunits (Fig. 5A,D; Methods). We did not observe any significant nucleotide-dependent changes in bending or twisting angles for FtsA and crenactin dimers (Fig. S8), likely because there was either little or no nucleotide dependence in monomer conformations of FtsA (Fig. 3) and crenactin (Fig. S7).

**Figure 5:**
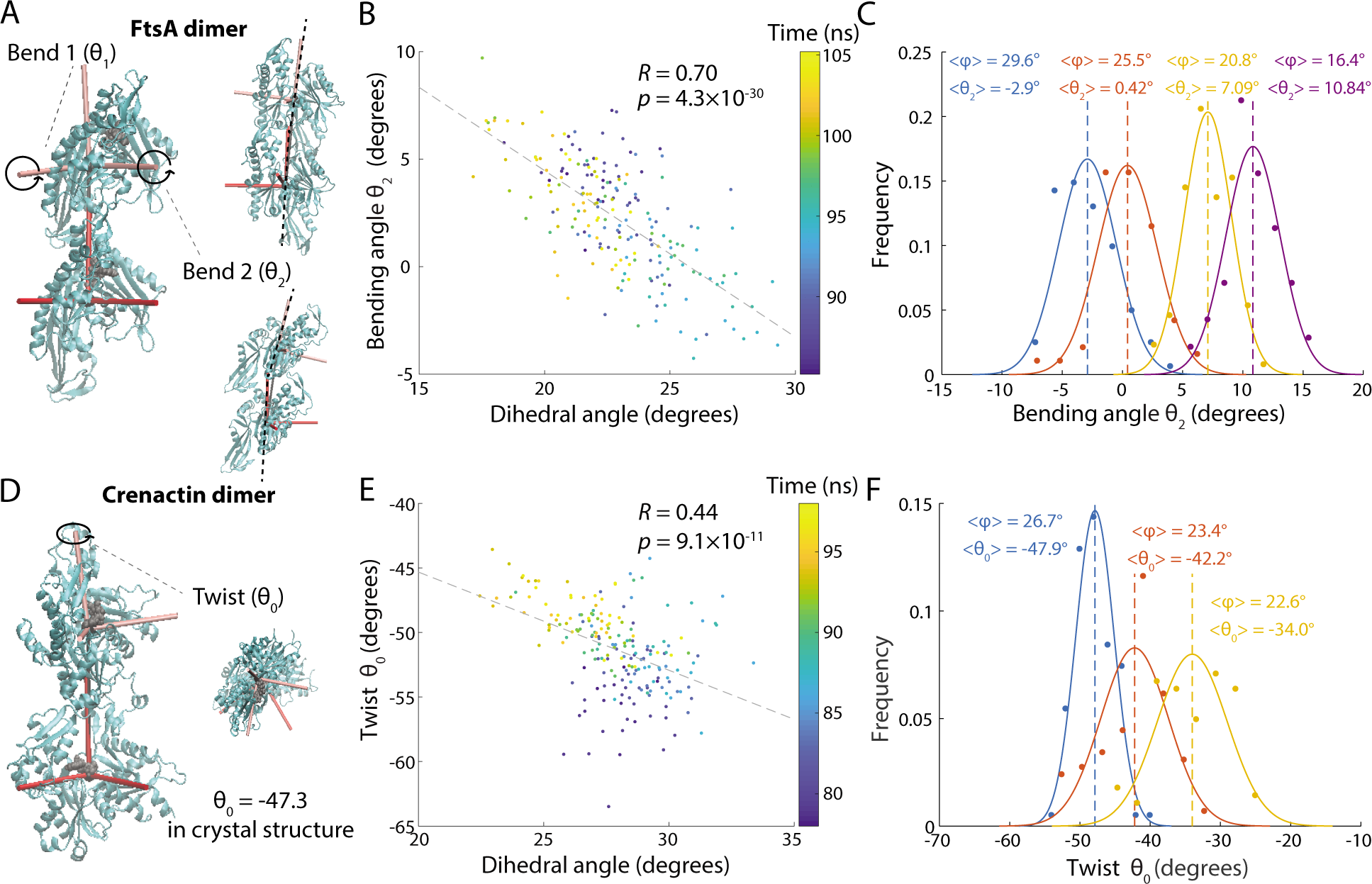
FtsA and crenactin filament bending and twisting are driven by changes to subunit dihedral angles. A) Illustration of the two possible axes for FtsA dimer bending. B) The dihedral angle of the bottom subunit in an FtsA-ATP dimer was highly correlated with bending angle *jθ*_*2*_ in all unconstrained simulations. C) Steering the dihedral angle *jθ*_*2*_ of the bottom subunit of an FtsA-ATP dimer from 16.4° to 29.6° caused systematic increases in the bending angle *θ*_*2*_. Curves are Gaussian fits to the data. Dashed lines are mean values. D) Illustration of the large degree of twist in a crenactin dimer. E) The dihedral angle *ϕ* of the bottom subunit in a crenactin-ATP dimer was highly correlated with dimer twist in unconstrained simulations. F) Steering the dihedral angle of the bottom subunit of a crenactin-ATP dimer from 22.6° to 26.7° caused a systematic increase in dimer twist. Curves are Gaussian fits to the data. Dashed lines are mean values.

Similar to MreB, the dihedral angle of the bottom subunit of an FtsA dimer was correlated with filament bending along the second bending axis (Fig. 5A,B). To test whether coupling between the dihedral angle and filament bending was direct, we steered the dihedral angle of the bottom subunit to average values of 16.4°, 20.8°, 25.5°, and 29.6° (measured over the last 20 ns of steered simulations; Fig. S9A). The resulting bending angles of the dimer shifted systematically with the dihedral angle (Fig. 5C), indicating that subunit dihedral changes drive bending of the FtsA filament. Interestingly, the bending angle flips from positive to negative (Fig. 5C); this flexibility could play a key role in regulating the transition of the division machinery from assembly to constriction.

In the crenactin filament crystal structure (PDB: 4CJ7), subunits have a large twisting angle of -47.3° (negative indicates right-handed filament); in our simulations, both ATP- and ADP-bound dimers equilibrated between -45° and -53° (Fig. S8D), suggesting that the large twisting angle is not a result of strained crystal contacts. By contrast to MreB and FtsA, the dihedral angle of the bottom subunit of crenactin was not correlated with filament bending, but rather with filament twist (Fig. 5D,E). To test causality, we steered the dihedral angle of the bottom subunit to 22.6°, 23.4°, and 26.7° (Fig. S9B), and observed progressive increases in twist magnitude (Fig. 5F). In sum, coupling of filament degrees of freedom to subunit conformational changes is generalizable across at least some bacterial actin-family members.

## Discussion

Through all-atom MD simulations of four actin-family proteins, we identified both conserved and specific dynamical behaviors across the actin family. First, we confirmed that the dihedral and opening angles between the centers-of-mass of the four subdomains represent the majority of conformational changes. In all simulated prokaryotic actins, the four subdomains exhibited high stability throughout the simulation, even as the whole protein changed conformation (Fig. 2E, 4B, S2, S7C). This analysis supports the model used by previous MD studies that measured dihedral and opening angles of actins [4, 13, 32], and provides a verified metric for future MD simulations of actin-family proteins.

Based on our findings, we propose a general model of the regulation of the structure of an actin-family filament in which the intra-subunit dihedral angle of a actin monomer regulates filament angles. The model suggests a mechanistic explanation for previous experimental results that have revealed variable filament structures for actin homologs. Electron microscopy of MreB, for instance, revealed straight filaments and arc-like filaments [6, 8]. Cryo-EM of crenactin filaments showed highly variable twists ranging from 32° to 56° [34]. Our simulations suggest that changes to bound-nucleotide state explain some of the variability in bend and twist for these dimers by tuning the dihedral angles of each subunit. Additionally, our finding that dihedral angle changes drive bending in FtsA and MreB but twisting in crenactin (Fig. 5, S6) indicate that the mechanism is not a trivial mechanical consequence of the four subdomain structure of actin homologs. Instead, the coupling between dihedral angle and key filament angles has likely been tuned for alternative filament behaviors over evolutionary time scales.

We observed distinct behaviors across actin homologs in terms of nucleotide dependence. MreB and ParM monomers exhibited distinct nucleotide-dependent states (Fig. 2A-D, 4A). These monomers have been shown to have ATPase activity [21, 35], suggesting that structural changes occur during the hydrolysis of ATP. Our results are also synergistic with efforts to translate the conformational variability of bacterial actin homologs for engineered purposes, including using ParM as a biosensor for ADP [36]. Numerous studies have attempted to detect ATPase activity in FtsA, but have found little or no activity [22, 37, 38]. Our simulations visualized distinct and reproducible nucleotide-dependent states (Fig. 3). Similar to our previous observation that the bending axis of an FtsA dimer rapidly changes upon release from crystal contacts [17], there is likely flexibility in the conformation of FtsA subunits that is masked in X-ray crystallography by symmetry requirements. For crenactin, we did not observe nucleotide dependence in monomer conformation in our simulations, all of which were carried out at 37 °C (Fig. S7). Crenactin has little ATPase activity at 37 °C, with maximum ATPase activity at 90 °C, which is far outside the temperature range for simulations with CHARMM force fields [12]. Thus, it remains to be seen whether crenactin behaves more like MreB/ParM or FtsA in its native environmental conditions of thermophilic temperatures. Hsp70, which forms a superfamily with actin based on a common fold, also exhibits nucleotide-dependent allostery [39], indicating that these intramonomeric changes may be general to a larger group of proteins. This basis for the large intramonomeric conformational changes in proteins such as MreB and ParM also suggests a strategy for the future design of proteins with similar flexibility and for the design of antibiotics that inhibit or disrupt these motions.

For prokaryotic actins, small perturbations in the protein’s environment can vastly impact structure. Many prokaryotic actins require binding proteins to confer their function *in vivo*, such as RodZ binding to MreB [40, 41]. Further, simulations of FtsA-FtsZ complexes could reveal why cell division relies upon the correct ratio of FtsA and FtsZ [42]. Crystal structures of FtsA-FtsZ complexes exist, but as we have shown with FtsA, crystal structures do not necessary capture the relevant physiological state, motivating the use of complementary techniques such as MD. In addition, genetic perturbations to prokaryotic actins can significantly impact cellular phenotypes. For example, mutations in MreB can have large effects on cell size and shape as well as MreB’s ability to sense curvature [43, 44]. Certain ParM mutations restrict the formation of helical filaments [45], and a variety of FtsA mutations restore viability after *zipA* deletion and alter cell shape [46–48]. Ultimately, crystallography, cryo-EM, *in vivo* light microscopy, and MD should prove a powerful combination for understanding and exploiting the numerous functions of cytoskeletal proteins.

## Methods

### MD simulations

All simulations (Table S1) were performed using the molecular dynamics package NAMD v. 2.10 [49] with the CHARMM36 force field [31], except where otherwise noted, including CMAP corrections [50],. Water molecules were described with the TIP3P model [51]. Long-range electrostatic forces were evaluated by means of the particle-mesh Ewald summation approach with a grid spacing of <1 Å. An integration time step of 2 fs was used [52]. Bonded terms and short-range, non-bonded terms were evaluated every time step, and long-range electrostatics were evaluated every other time step. Constant temperature *(T* = 310 K) was maintained using Langevin dynamics [53], with a damping coefficient of 1.0 ps^−1^. A constant pressure of 1 atm was enforced using the Langevin piston algorithm [54] with a decay period of 200 fs and a time constant of 50 fs. Setup, analysis, and rendering of the simulation systems were performed with the software VMD v. 1.9.2 [55]. Steering of the dihedral angle and of distances between residues was achieved by introducing collective forces to constrain angles and distances to defined values through the collective variable functionality of NAMD [49].

### Simulated systems

MD simulations performed in this study are described in Table S1. For simulated systems initialized from a MreB crystal structure, the crystallographic structure of *T. maratima* MreB bound to AMP-PMP (PDB ID: 1JCG) [4] was used; for FtsA, the crystallographic structure of *T. maratima* FtsA bound to ATP gamma A (PDB ID: 4A2B) was used; for ParM, the crystallographic structure of *E. coli* ParM (PDB ID: 1MWM) bound to ADP was used; for crenactin, the crystallographic structure of *P. calidifontis* crenactin bound to ADP (PDB ID: 4CJ7) [12] was used. The bound nucleotide was replaced by both ATP and ADP for all simulated systems, and Mg^2+^-chelating ions were added for stability. Water and neutralizing ions were added around each monomer or dimer, resulting in final simulation sizes of up to 157,000 atoms. All unconstrained simulations were run for 54-134 ns. All steered simulations were run until equilibrium was reached. For mean values and distributions of measurements, only the last 30 ns of unconstrained simulations or the last 20 ns of steered simulations were used. To ensure simulations had reached equilibrium, measurement distributions were fit to a Gaussian.

### Analysis of dihedral and opening angles

The centers-of-mass of the four subdomains of each protein were obtained using VMD. For each time step, we calculated one opening angle from the dot product between the vector defined by the centers-of-mass of subdomains IIA and IIB and the vector defined by the centers-of-mass of subdomains IA and IB (or IC for FtsA). Similarly, we calculated a second opening angle from the dot products between the vectors defined by the centers-of-mass of subdomains IA and IB and of subdomains IIA and IA. The opening angles we report are the average of these two opening angles. The dihedral angle was defined as the angle between the vector normal to a plane defined by subdomains IA, IB, and IIA and the vector normal to a plane defined by subdomains IIB, IIA, and IA. Subdomain definitions for each protein are provided in Table S2.

### Analysis of bending and twisting angles

At each time step of a dimer simulation, the coordinate system of the bottom and top monomers was defined using three unit vectors {**d**1, **d**2, **d**3}. **d**_1_ approximately aligns to the center-of-mass between the two subunits, and **d**3 is defined to be zero at the start of the simulation. Rotation around **d**i represents twist between the bottom and top subunits. Since **d**3 is defined to be zero at the start of the simulation, **d**2 represents the ideal bending axis. **d**3 represents bending in a direction orthogonal to **d**2.

## Supplementary Tables

**Table S1:**
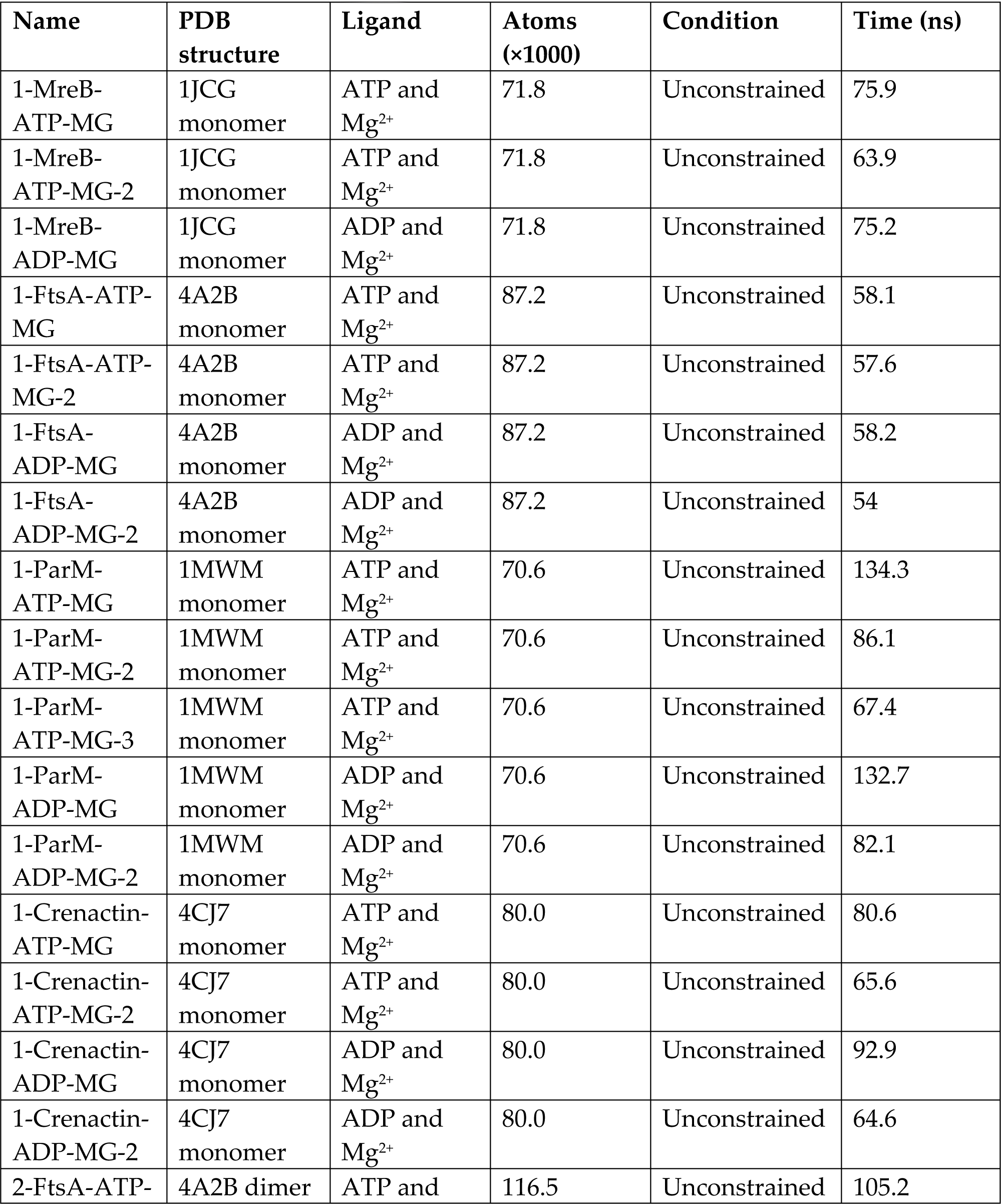
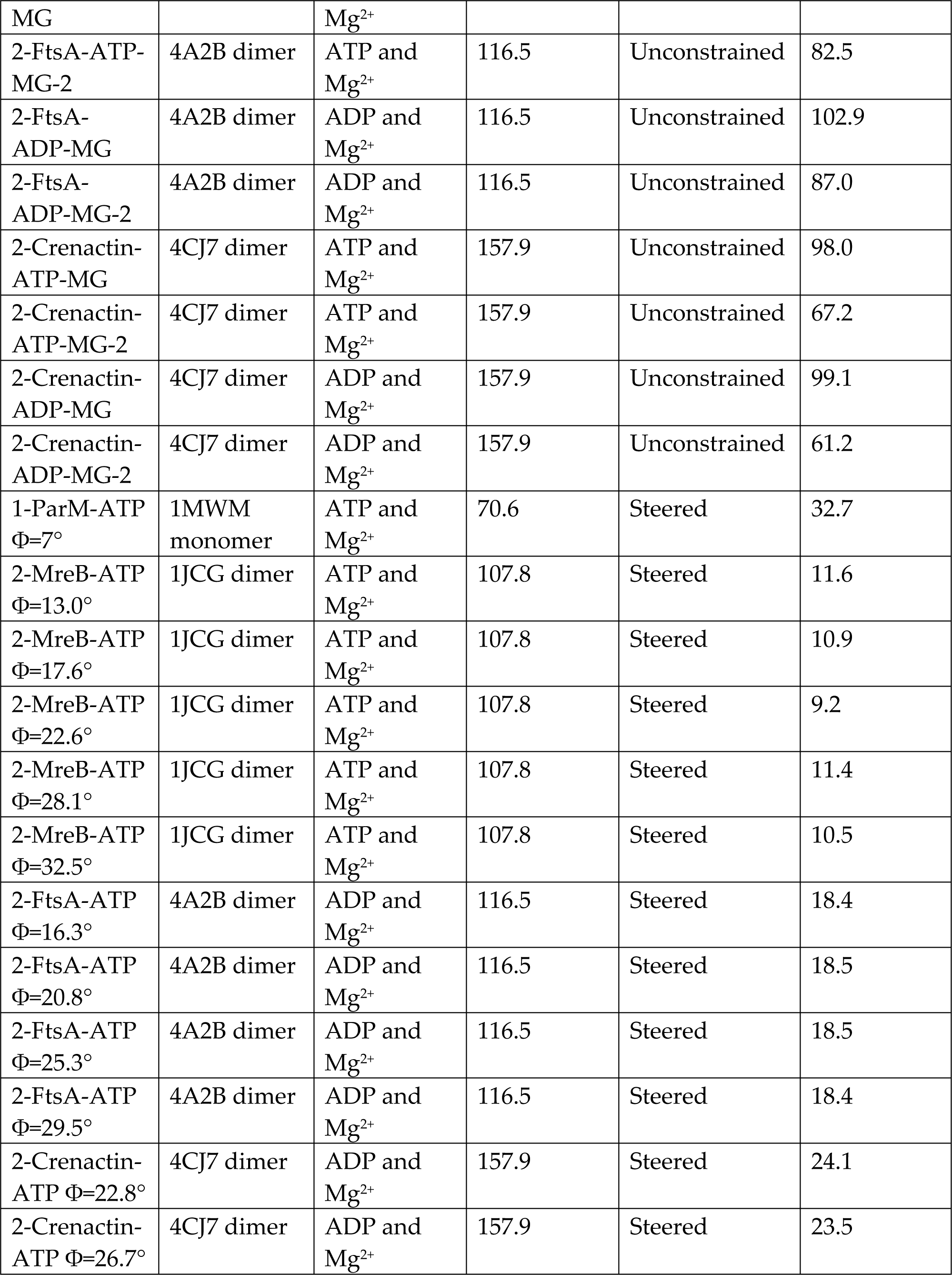
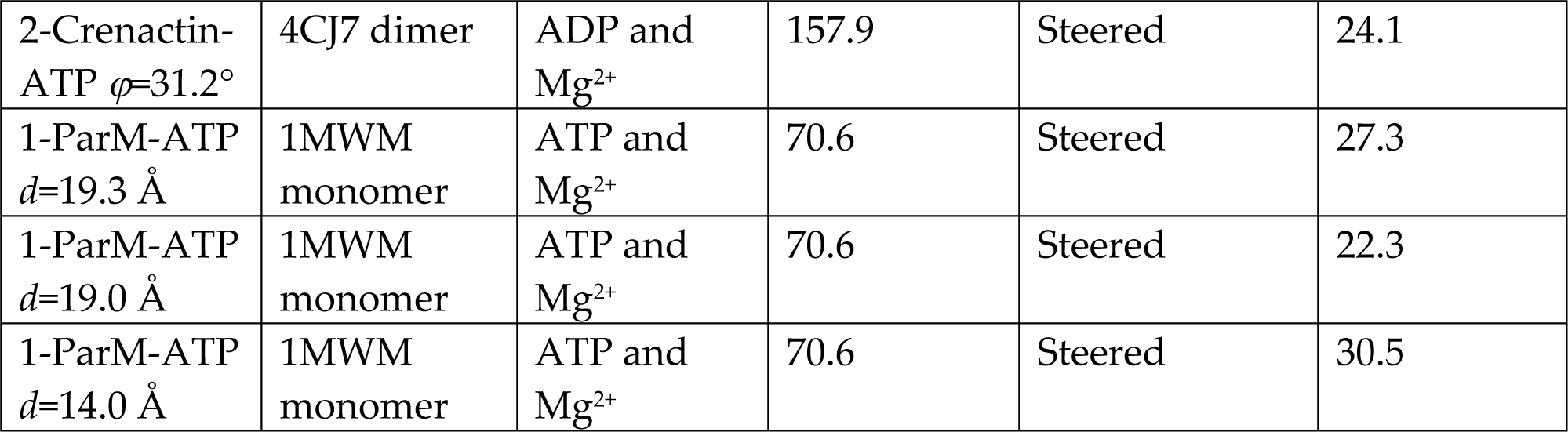
MD simulations in this study.

**Table S2:** Subdomain definitions by residue numbers.

| Protein | Structure | IA | IB | IIA | IIB | IC |
| --- | --- | --- | --- | --- | --- | --- |
| MreB | 1JCG monomer | 1-138; 315-336 | 30-72 | 139-175;<br>251-314 | 176-249 | N/A |
| FtsA | 4A2B monomer | 1-86; 167-198;<br>360-392 | N/A | 199-234;<br>305-359 | 235-304 | 87-166 |
| ParM | 1MWM monomer | 1-29, 68-159;<br>306-320 | 30-67;<br>130-137 | 160-198;<br>255-305 | 199-254 | N/A |
| Crenactin | 4CJ7 monomer | 1-38; 119-173;<br>399-432 | 39-118 | 174-207;<br>300-398 | 208-194 | N/A |

## Acknowledgments

We thank Jen Hsin and the Huang lab for helpful discussions. Funding was provided by NSF CAREER Award MCB-1149328, the Stanford Center for Systems Biology under Grant P50-GM107615, and the Allen Discovery Center at Stanford on Systems Modeling of Infection (to K.C.H); an Agilent Fellowship and a Stanford Interdisciplinary Graduate Fellowship (to H.S.); and a Stanford Graduate Fellowship and a Gerald J. Lieberman Fellowship (to A.C.). N.N. was supported by the Stanford Bioengineering Research Experience for Undergraduates program. K.C.H. is a Chan Zuckerberg Investigator. All simulations were performed with computer time provided by the Extreme Science and Engineering Discovery Environment (XSEDE), which is supported by National Science Foundation grant number 0CI-1053575, with allocation number TG-MCB110056 (to K.C.H.). This work was also supported in part by the National Science Foundation under grant PHYS-1066293 and the hospitality of the Aspen Center for Physics.

